# All talk? Left temporal alpha oscillations are not specific to verbal-analytical processing during conscious motor control

**DOI:** 10.1101/851956

**Authors:** Johnny V. V. Parr, Germano Gallicchio, Neil R. Harrison, Ann-Kathrin Johnen, Greg Wood

## Abstract

The present study tested the validity of inferring verbal-analytic motor processing from EEG left-temporal alpha activity. Participants (*n* = 20) reached for and transport a jar under three conditions: one control condition and two self-talk conditions aimed at eliciting either task-unrelated verbal processing or task-related conscious control, while 32-channel EEG and kinematics were recorded. Compared to the control condition, both self-talk conditions elicited greater self-reported levels of verbal processing, but only the task-related self-talk condition was accompanied by greater left temporal activity (i.e., EEG alpha power decreased) during movement production. However, this increase was not localised to the left temporal region but was rather evident over all scalp regions examined, suggesting an interpretation more consistent with diminished neural efficiency. No effects for left temporal-frontal (T7-Fz) connectivity were detected across conditions. Our results failed to endorse left-temporal EEG alpha activity as valid index of verbal-analytic processing during motor tasks.

## 1. Introduction

The progression from beginner to skilled motor performance is characterised by an attenuation of energy expenditure as the expression of greater metabolic efficiency (Hatfield, 2018; Hatfield & Hillman, 2001). Such adaptations are not only observed as decreased somatic activity (e.g., reduced muscular activation), but also as decreased mental activity (e.g., reduced regional activation in the brain). By using neuroimaging techniques, such as electroencephalography (EEG), researchers have provided evidence that practice of a motor skill induces changes in the cerebral cortex consistent with the concept of “neural efficiency”.

A cortical region that is often deemed to be implicated in the attainment of greater neural efficiency is the left temporal region. By recording the magnitude of EEG oscillatory activity within the 8-12 Hz frequency range, an inverse marker of neuronal activity termed “alpha power” (Klimesch, 2012), researchers have observed diminished activity in the left temporal region as a function of expertise (Haufler, Spalding, Santa Maria, & Hatfield, 2000; Janelle et al., 2000), training (Kerick, Douglass, & Hatfield, 2004), and performance (Gallicchio, Cooke, & Ring, 2017). As the left temporal region includes structures implicated in language processing (e.g., Broca’s area and Wernicke’s area), the abovementioned findings have been interpreted as evidence that expert performance is less reliant on declarative verbal-analytic processes that characterise the conscious motor control of novices (e.g., Fitts & Posner, 1967).

In addition to regional activation, cortico-cortical networking has been examined to reveal the interaction across various cortical regions. For example, phase-based measures of alpha connectivity between the left temporal (T7) and the frontal premotor region (Fz) have been examined to assess the influence of declarative verbal processing on the production of movement. Alpha connectivity reflects the synchronicity in the inhibition profiles of two regions, with greater alpha connectivity suggesting consistently similar inhibition (functional communication) and lower alpha connectivity suggesting distinct inhibition profiles (regional independence). In line with the acquisition of greater automaticity with training, researchers have shown that T7-Fz upper-alpha (~10-12 Hz) connectivity decreases with increasing skill (Deeny, Hillman, Janelle, & Hatfield, 2003; Gallicchio, Cooke, & Ring, 2016; Gallicchio et al., 2017; Kerick et al., 2004). This is suggestive of a gradual disconnect between the verbal-analytical and premotor regions as individuals consolidate movement patterns and efficiently organise task-related neural networks free from conscious control. Further work has shown that T7-Fz upper-alpha connectivity increases when directing participants to exert conscious movement control (Ellmers et al., 2016) and is greater for novices who undertake explicit motor learning (high exposure to declarative knowledge) compared to those who undertake implicit motor learning (low exposure to verbal-analytic rules) (Parr, Vine, Wilson, Harrison, & Wood, 2019; Zhu, Poolton, Wilson, Hu, et al., 2011). Finally, participants who report a strong propensity to consciously monitor and control their movements also display increased T7-Fz connectivity compared with those with a lower propensity (Zhu, Poolton, Wilson, Maxwell, & Masters, 2011).

Taken together, these findings suggest that skilled, autonomous, expert-like motor performance is associated with decreased left temporal involvement in the production of movement. Conversely, less skilled, conscious, and novice-like motor performance is associated with increased left temporal involvement in the production of movement. As the left temporal region is associated with language, these findings fit well with classic models of motor learning that describe an early reliance on declarative verbal knowledge to guide performance that becomes subsequently “proceduralised” into a non-verbal memory units as skill progresses (Fitts & Posner, 1967). Left temporal alpha oscillations would therefore appear a useful yardstick for verbal-analytical processes during motor control.

Crucially, implicit within this proposition is the assumption that EEG activity recorded over left temporal sites (i.e., T7) during motor execution is uniquely representative of the underlying cortical structures associated with language (Cooke, 2013). However, this may not be the case for two reasons: first, unless a one-to-one mapping can be demonstrated, it is not deductively valid to infer a particular cognitive process from the activation of a particular brain region. This practice, termed “reverse inference”, reasons backwards from the presence of brain activation to the engagement of a particular cognitive function, and is limited by non-selectivity of activation in the region of interest (Poldrack, 2006). For example, structures across the left temporal region are implicated not only in language processing but also in auditory processing (Tiihonen et al., 1991) and the integration of visual and auditory information (Beauchamp, Lee, Argall, & Martin, 2004). Second, the EEG has a relatively low spatial resolution due to the propagation of electrical fields across tissues – a phenomenon referred to as “volume conduction” – meaning that the activity recorded from a certain channel can be significantly influenced not just by a local source but also by large distal sources (Cohen, 2015).

Consequently, it is unclear the extent to which EEG alpha activity recorded over left temporal sites may reflect a broader range of processes associated with unskilled and conscious motor control beyond that of language, such as general demands on attention and motor effort. This issue is corroborated by research showing that, compared to experts, novices display a global decrease in alpha power that is not specific to the left temporal region (Babiloni et al., 2010; Baumeister, Reinecke, Liesen, & Weiss, 2008; Del Percio et al., 2009; Janelle et al., 2000; Parr et al., 2019). It is therefore plausible that differences in left temporal alpha during motor control are more reflective of general reductions in neural efficiency rather than uniquely reflective of verbal processing.

The aim of this study was to manipulate the content of verbal-analytical processing to explore how language processing affects measures of regional alpha power and connectivity during the planning and production of reaching and grasping movements. Specifically, we compared task-related and task-unrelated self-talk with uninstructed, natural performance. Whilst both types of self-talk were administered to increase verbal processing during motor performance, only task-related (declarative) self-talk was designed with the intention of interfering with movement automaticity and increasing conscious motor processing (Masters, 1992). If left temporal alpha activity recorded during motor execution uniquely reflects verbal-analytical processing, then we would expect *both* conditions that encourage self-talk (task-related and task-unrelated) to increase activity in the left temporal region (decreased lower alpha power). However, if self-talk induces changes in regions other than left temporal, this would support the thesis that the association between left temporal activity and language processing during motor performance can be attributed to a spatially broader phenomenon, possibly consistent with decreased neural efficiency. Finally, if left temporal alpha activity were related to the functional connectivity between the left temporal and premotor regions (T7-Fz) during movement production, then we would only expect increased connectivity in the task-related self-talk condition.

## 2. Methods

### 2.1 Participants

Twenty self-reported right-handed participants took part in the study (11 females, 9 males, age: *M* = 26.38, *SD* = 6.19 years). All participants gave written consent and the procedures were approved by our institutional ethics committee.

### 2.2 Experimental task

The task required participants to sequentially reach for and then transport a glass jar from a home position to a target position on a desk (Figure 1). Prior to each trial, participants placed their right hand on the start location positioned at the desk edge approximately 30cm away from the jar home position. The trial started with the onset of an auditory tone (S1) indicating the start of a 2-second preparation period. During this period, participants were instructed to stay still, maintain their hand on the start location, focus their gaze on the jar, and avoid eye blinks and body movement to minimise EEG data artefacts. At the end of this 2-second period, a second auditory tone (S2) acted as a “go” signal to initiate the task. After placing the jar on the target position, participants returned their hand to the start location and the researcher prepared the workspace for the following trial. Participants were instructed to perform at a steady speed that felt comfortable and natural. In order to decrease the repetitive nature of the task and increase task engagement, the weight of the jar was randomly varied at each trial among five options (250, 350, 450, 550, 650 g). Auditory tones (each 0.67 seconds long) were controlled through E-Prime 2.0 (Psychology Software Tools, Inc., Pittsburgh, PA, USA).

**Figure 1.**
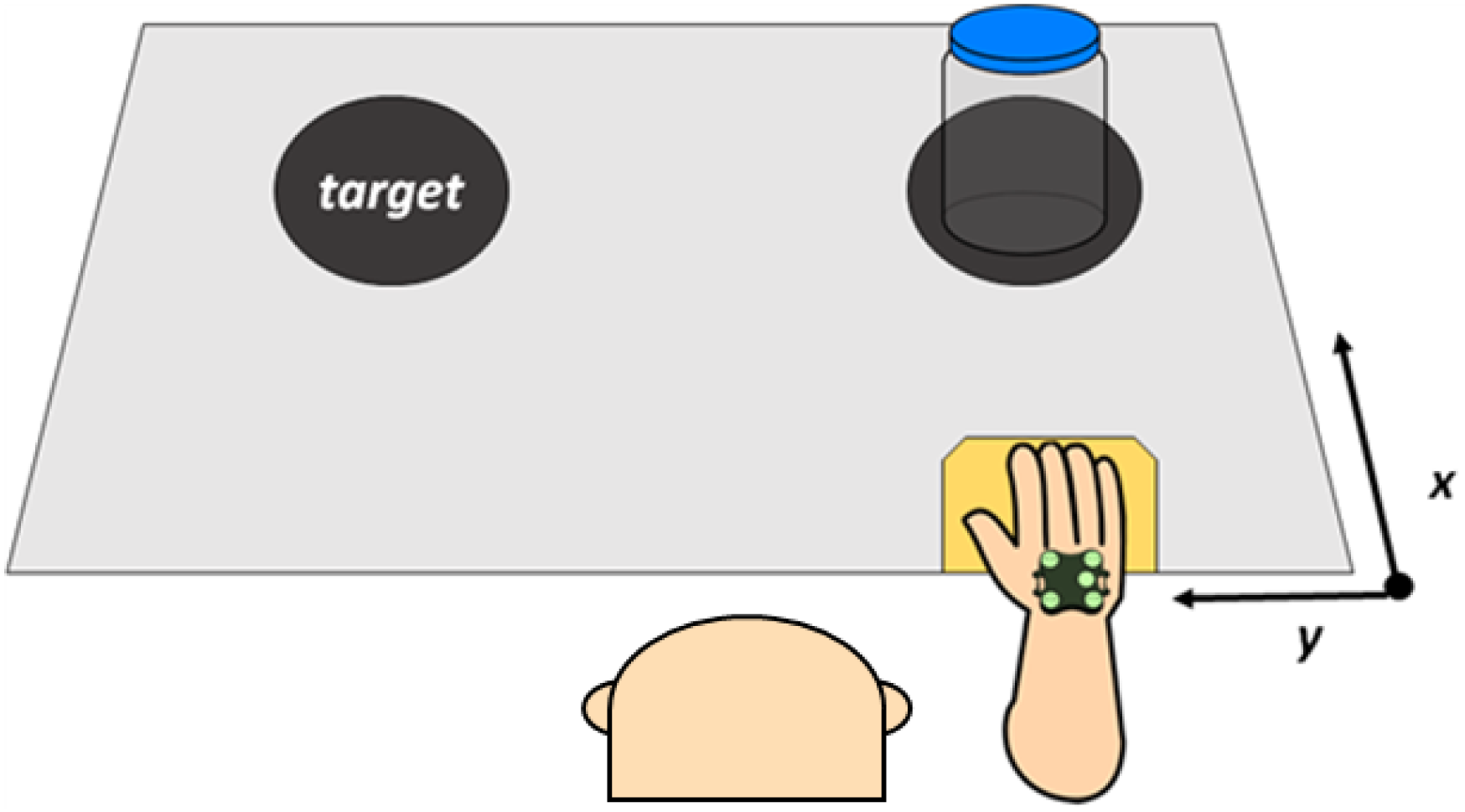
Visual representation of the experimental set-up. Participants began each trial with their right hand on the start position (in yellow). A first auditory tone (S1) indicated the start of the 2-second preparation period, during which participants maintained their hand on the start position and their gaze on the jar. Following a second auditory tone (S2) participants reached for the jar and placed it on the target location.

### 2.3 Procedure

Participants attended one 2-hour session. After briefing, participants were seated at a distance that enabled them to reach the jar home and target positions at arms-length. Once instrumented for EEG and kinematic recording, participants completed a 1-minute eyes-closed resting baseline and then completed 10 practice trials to enable familiarise the task. Participants then performed 40 trials in the *control condition*, which consisted of reaching and transporting the jar with no self-talk instructions. Participants then completed manipulation checks at the end of the 40 trials assessing task difficulty, mental load, conscious control, and self-talk (see below for details).

Then, the experiment was performed under two *self-talk conditions* – one task-related and one task-unrelated – each composed of 40 trials. These trials were performed in eight blocks of ten in an interleaved manner (e.g., 10 x task-related, 10 x task-unrelated, 10 x task-related etc.), with the starting condition counterbalanced across participants. For the task-related condition, self-talk instructions regarded the control of movements to encourage movement conscious processing: “*Keep elbow below the wrist*”, “*Keep palm 5(cm) from table*”, “*Keep thumb below the index*”, “*Keep wrist flexed at 90 degrees*”. For the task-unrelated condition, self-talk instructions included a collection of well-known nursery rhymes that were not related to movement: “*Jack and Jill went up the hill”, “Twinkle twinkle little star”, “Mary had a little lamb”, “Humpty dumpty sat on a wall”*. The to-be-rehearsed self-talk instruction varied in a random manner for each block of 10 trials. Participants were required to silently rehearse these instructions throughout the entirety of each trial (i.e., during both movement preparation and execution phases) to the best of their ability. For both self-talk conditions, participants were asked which word of the given phrase they finished on when placing each jar. This word was written down in view of each participant to act as a manipulation check and to encourage adherence to self-talk instructions. Adherence to the task-related self-talk condition was further encouraged by positioning electrodes on the right upper-limb and workspace prior to each block of 10 trials to convince participants that their adherence to the task-related instructions would be quantitatively evaluated (Appendix S1). After each of the ten blocks of trials, participants again rated task difficulty, mental load, conscious control, and self-talk (see below for details).

### 2.4 EEG

Thirty-two active EEG electrodes were positioned on the scalp at locations Fp1, Fp2, AF3, AF4, F7, F3, Fz, F4, F8, FC5, FC1, FC2, FC6, T7 (T3), C3, Cz, C4, T8 (T4), CP5, CP1, CP2, CP6, P7, P3, Pz, P4, P8, PO3, PO4, O1, Oz and O2 of the 10-20 system (Jasper, 1958)^1^. Common mode sense (CMS) and driven right leg (DRL) electrodes were used to enhance the common mode rejection ratio of the signal. The signal was amplified and digitized at 512 Hz using the ActiveTwo recording system (Biosemi, The Netherlands). A digital trigger was automatically sent from E-Prime to the Biosemi system via parallel communication at the onset of each S2 (“go” signal). All EEG signals were re-referenced to the average of all channels and 1-35 Hz band-pass filtered (FIR [finite infinite response]). Epochs were cut from −2.25 to +2.25 s relative to the onset of S2. Epochs were then visually inspected so those showing movement artefact could be rejected (percentage of trials rejected: *M* = 13.33, *SD* = 5.89 %). No bad channels were identified. Data were then subjected to Independent Component Analysis (Runica Infomax algorithm; Makeig, Bell, Jung & Sejnowski, 1996) to identify and remove components accounting for blinks, eye movements and other non-neural activity (components removed: *M* = 5.70, *SD* = 3.05). These processing steps were performed using EEGLAB functions (Delorme & Makeig, 2004) and bespoke MATLAB scripts.

### 2.5 Kinematic data

Noraxon MyoMotion (Scottsdale, AZ, USA) motion analysis system was employed to analyse kinematic characteristics of the task performance. A single MyoMotion inertial measurement unit (IMU) was placed on the dorsal side of the right hand according to the rigid-body model defined in the Noraxon MR3 software. Calibration was performed using the upright standing position, though the use of a single IMU wavered the need for a zero/neutral angle in the measured joints. The sampling frequency for the inertial sensor was set at 100Hz. To identify movement onset (S2) for each trial, a tripod-mounted webcam (30Hz) was synchronised using a light-emitting diode (LED) and Noraxon software to enable digital triggers to be applied to the data offline following visual inspection of the recorded video. Data were then smoothed using a 10-point (corresponding to 100ms) moving average before being epoched from −2.25 to + 2.25 s relative to S2 (“go” signal) for each trial.

### 2.6 Measures

#### 2.6.1 Manipulation check

Manipulation checks were administered throughout testing to confirm that our experimental conditions were having their desired effect. Using 7-point Likert scales, participants were asked to self-report levels of task difficulty, mental effort, movement awareness, movement control, self-talk frequency, and self-talk intensity.

#### 2.6.2 Movement time

Movement time was measured as the time (in seconds) elapsed between S2 (i.e., “go” signal) and jar placement on the target location. To obtain this, the researcher provided a keyboard response upon jar placement, enabling movement time to be extracted from E-Prime.

#### 2.6.3 Peak acceleration

The early characteristics of reaching and grasping movements can provide useful insights into the extent to which movements are performed under predictive or online control. For example, increased limb acceleration indicates more ballistic and pre-planned movement, whereas decreased limb acceleration would suggest a more conservative approach with greater reliance on feedback to supervise ongoing action (Desmurget & Grafton, 2000). Peak acceleration along the *x*- and *y*-axes of movement was therefore measured for the reach and transport phases of our task to provide kinematic evidence of how our self-talk instructions interfered with movement automaticity. Specifically, peak reach acceleration was taken as the peak value recorded between 0 and +500 ms relative to S2 along the *x*-axis of movement. Peak transport acceleration was taken as the peak value recorded between 500 and 1500 ms relative to S2 along the *y*-axis of movement. Separate axes were chosen for each phase as they each capture the primary direction of movement. Time windows were selected upon visual inspection of data from which clear peaks for each phase could be easily detected (see Figure 4).

#### 2.6.4 Alpha power

Time-frequency decomposition of the epoched EEG signals was performed for each trial through short-time Fast Fourier Transform (FFT) conducted on 37 overlapping windows, each of 0.5 s (87.5% overlap), with central points ranging from −2 to +2 s. Prior to FFT, data points in each window were Hanning-tapered and zero-padded to reach 4 s. This procedure generated complex-valued coefficients in the time-frequency plane with a precision of 111ms and 0.25 Hz, separately for each channel and trial. Alpha power was then computed as the squared amplitude within each participant’s individual alpha frequency (IAF) band, identified using both the ‘peak frequency’ and ‘centre of gravity’ (CoG) methods (Corcoran, Alday, Schlesewsky, & Bornkessel-Schlesewsky, 2018). To do this, we first attempted to identify a visible peak (between approximately 7 – 15Hz) from the mean EEG spectrum over occipito-parietal channels, recorded from the 1-minute eyes-closed baseline. When a single peak was not present (40% of participants), IAF was instead taken as the power spectra weighted mean (CoG) between 7 and 15 Hz (Appendix S2). Subsequently, the alpha frequency band was denoted as IAF-2 to IAF+2. For one participant, baseline data was corrupted, resulting in alpha being denoted as the typical 8 – 12 Hz range. As no neutral baseline could be identified, non-normal distributions and inter-individual differences were dealt with by employing a median-scaled log transformation. This transformation was implemented by scaling all alpha power values for each participant (across all channels, trials, time-windows, and conditions) by the median alpha power value within that participant, before then employing a 10·log10 transformation. Power was then averaged across overlapping time windows and across trials to provide mean values for each channel, separately from −2s to −1s (*t^−2^*), −1s to 0s (*t^−1^*), 0s to 1s (*t^+1^*), and 1s to 2s (*t^−2^*). To examine regional effects, seven regions of interest (ROI) were identified based on inspection of topographical maps: frontal (F3, Fz, F4), left temporal (F7, FC5, T7), left central (FC1, C3, CP1), right central (FC2, C3, CP2), right temporal (F8, FC6, T8), parietal (P3, Pz, P4) and occipital (O1, Oz, O2). Values within each ROI were averaged. Signal processing was performed in MATLAB.

#### 2.6.5 Alpha connectivity

Functional connectivity was computed as the inter-site phase clustering (ISPC) over time (Cohen, 2014; Lachaux, Rodriguez, Martinerie, & Varela, 1999). ISPC measures the phase lag consistency across time between two channels independently from their power and reflects functional connectivity between the oscillatory activity of two underlying cortical regions, with values ranging from 0 (no connectivity) to 1 (perfect connectivity). ISPC was calculated for each trial using bespoke Matlab scripts as, 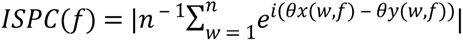, where *i* is the imaginary operator; *θx* and *θy* are the phase angles of the recorded signal at two different scalp locations at FFT time window *w* and frequency *f*; *e*^*i*(*θx*(*w*,*f*) ‒ *θy*(*w*,*f*))^ denotes a complex vector with magnitude 1 and angle 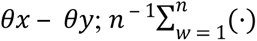 denotes averaging across the overlapping FFT time windows within each predefined epoch (*t^−2^, t^−1^, t^+1^, t^+2^*); and |·| is the module of the average vector. Following previous research and our a priori hypotheses, we focused on left temporal (T7) – frontal (Fz) ISPC in the upper-alpha (IAF to IAF+2) frequency band for our statistical analyses. Right temporal (T8) – frontal (Fz) upper-alpha connectivity was also analysed to verify the extent to which connectivity effects were localised to a given hemisphere. No baselines were used. Instead, to normalise their density distributions, ISPC were Fisher *Z* transformed (inverse hyperbolic tangent); values could then range from 0 to ∞. Values were then averaged over trials and frequencies within the upper-alpha range (IAF to IAF+2 Hz) to yield estimates of alpha connectivity separately for each condition.

### 2.7 Statistical analyses

For self-report data, a one-way repeated measures ANOVA was performed to assess the effect of Condition (control, task-related, task-unrelated) on each of the six items (difficulty, mental effort, movement awareness, movement control, self-talk frequency, self-talk intensity). One-way repeated measures ANOVAs were also performed to assess the effect of Condition on movement time, peak reach acceleration and peak transport acceleration.

A 3 Condition x 4 Epoch (*t^−2^*, *t^−1^*, *t^+1^*, *t^+2^*) repeated measures ANOVA on the averaged alpha power over channels T7, F7 and FC5 was performed to specifically test the prediction that left temporal alpha power can be used to infer verbal processing. Following this, a 3 Condition x 4 Epoch x 7 ROI (left temporal, frontal, left central, right central, parietal, right temporal, occipital) repeated measures ANOVA was conducted to evaluate the presence of wider regional effects of alpha power. Finally, a 3 Condition x 4 Epoch x 2 Hemisphere (T7-Fz, T8-Fz) repeated measures ANOVA was conducted to assess time-varying changes in left and right temporal-frontal alpha connectivity across Conditions. By comparing the 2 s preceding movement with the 2 s following movement, we can determine whether effects are specific to the preparation or execution of motor performance.

For all ANOVAs, Greenhouse-Geisser corrections were applied when sphericity was violated and effect sizes were calculated using partial eta squared (np2). All post hoc pairwise comparisons were adjusted using Bonferroni corrections to counteract the problem of multiple comparisons.

## Results

### 3.1 Manipulation checks

Results from each one-way repeated measures ANOVA showed a significant main effect of Condition for task difficulty, *F*(2, 36) = 66.04, *p* < .001, np2 = .786, mental effort, *F*(2, 36) = 49.37, *p* < .001, np2 = .733, movement awareness, *F*(2, 36) = 57.56, *p* < .001, np2 = .762, conscious control, *F*(2, 36) = 77.08, *p* < .001, np2 = .811, self-talk frequency, *F*(2, 36) = 27.37, *p* < .001, np2 = .603, and self-talk intensity, *F*(2, 36) = 14.739, *p* < .001, np2 = .450. Post-hoc pairwise comparisons showed significantly higher levels of self-talk frequency (*p* < .001) and self-talk intensity (*p* < .001) during the task-related and task-unrelated self-talk conditions compared to the control condition, confirming the intended manipulation of verbal processing. Task difficulty, mental effort, movement awareness, and conscious control were rated significantly higher in the task-related condition compared to the task-unrelated and control conditions (*p* < .001), but the task-unrelated condition as significantly more difficult than the control condition (*p* = .013).

**Table 1.** Means and standard deviations of self-report and kinematic data, together with the results from each one-way repeated measures ANOVA.

|  | **Control**<br>**M (SD)** | **Task-**<br>**related**<br>**M (SD)** | **Task-**<br>**unrelated**<br>**M (SD)** | ***F*** | ***np*<sup>2</sup>** |
| --- | --- | --- | --- | --- | --- |
| Difficulty | 1.05 (0.23) | 3.33 (1.09) | 1.50 (0.62) | 66.04* | .79 |
| Mental effort | 1.47 (0.96) | 4.14 (1.00) | 2.07 (1.00) | 49.37* | .73 |
| Movement<br>awareness | 3.16 (1.30) | 5.74 (1.09) | 2.88 (0.81) | 57.56* | .76 |
| Movement<br>control | 2.47 (1.07) | 5.71 (0.85) | 2.39 (1.08) | 77.08* | .81 |
| Self-talk<br>frequency | 4.16 (1.53) | 6.54 (0.56) | 6.43 (0.69) | 27.37* | .60 |
| Self-talk<br>intensity | 3.89 (1.79) | 5.92 (0.88) | 5.81 (0.86) | 14.74* | .45 |
| Reach<br>(peak m/s<sup>2</sup>) | 2.20 (0.52) | 1.49 (0.40) | 2.04 (0.47) | 34.38* | .86 |
| Transport<br>(peak m/s<sup>2</sup>) | 1.70 (0.37) | 1.01 (0.19) | 1.55 (0.37) | 31.21* | .68 |
| Movement time<br>(sec) | 1.64 (0.21) | 2.66 (0.51) | 1.86 (0.34) | 74.56* | .80 |

\* indicates a significant main effect of Condition ( $p < .001$ ).

### 2.8 Movement time

The one-way repeated measures ANOVA showed a significant main effect of Condition, *F*(2, 38) = 74.56, *p* < .001, np2 = .797. Post-hoc pairwise comparisons revealed participants performed slowest during the task-related condition (*p* < .001), and fastest during the control condition (*ps* = .005).

### 2.9 Peak acceleration

A one-way repeated measures ANOVA showed a significant main effect of Condition for both the reach, *F*(2, 32) = 34.38, *p* < .001, np2 = .862, and transport, *F*(2, 30) = 31.21, *p* < .001, np2 =.675, phases of the task. Pairwise comparisons showed peak acceleration to be lower for the task-related condition, compared to control and task-unrelated conditions for both reach (*p* < .001) and transport (*p* < .001) task phases (Figure 2).

**Figure 2.**
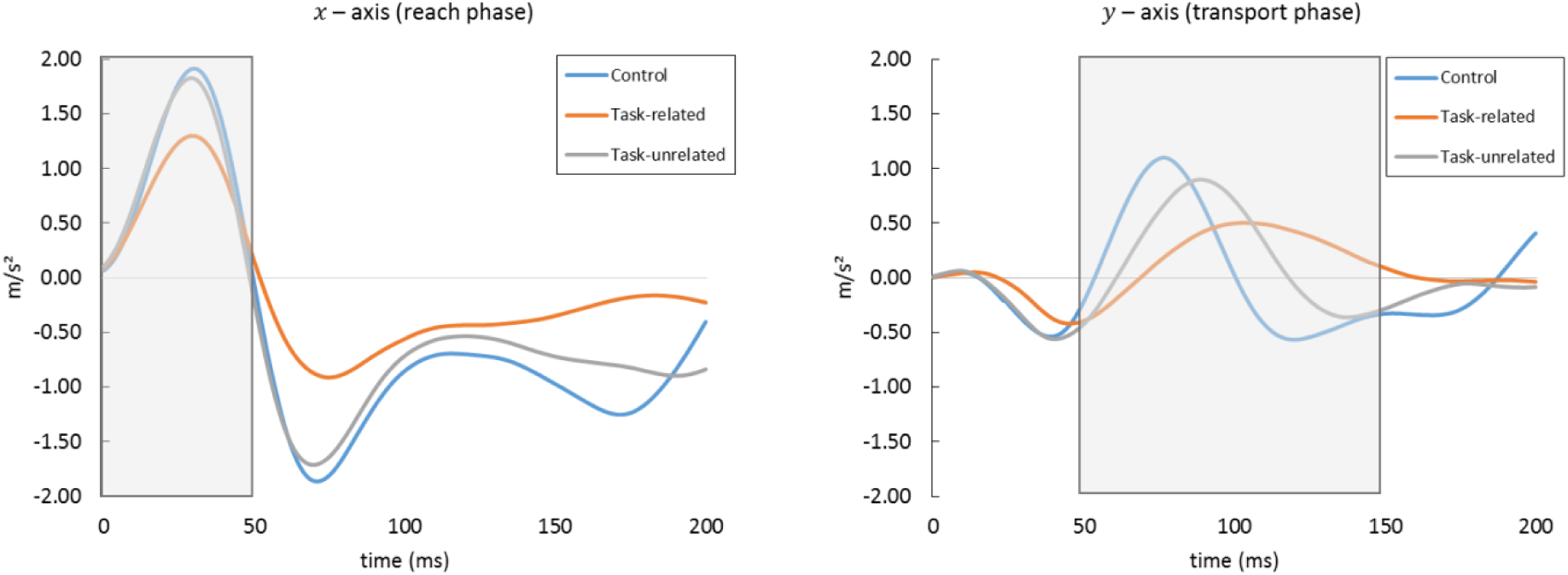
Mean acceleration profiles over time for each condition. Shaded grey areas depict the time window in which peak reach acceleration (left) and peak transport acceleration (right) were identified.

### 2.10 Alpha power

#### 2.10.1 Left temporal alpha

The 3 (Condition) x 4 (Epoch) repeated measures ANOVA conducted on the averaged activity recorded from channels T7, F7 and FC5 yielded a marginally significant main effect of Epoch, *F*(1.54, 29.26) = 3.273, *p* = .064, np2 = .147, and a significant main effect of Condition, *F*(2, 38) = 7.031, *p* = .003, np2 = .270. The Condition x Epoch interaction was non-significant, *F*(6, 114) = 0.294, *p* = .938, np2 = .015. Post-hoc pairwise comparisons to interrogate the main effects revealed that left-temporal alpha power decreased from *t^+1^* to *t^+2^* (*p* = .001). They also revealed lower left-temporal alpha power in the task-related self-talk condition, compared to the control (*p* = .042) and task-unrelated self-talk (*p* = .002) conditions.

#### 2.10.2 Regional alpha

These analyses were conducted to examine topographical effects involving multiple regions of interest covering the major areas of the cerebral cortex, including, but not limited to, the left temporal region. Results from a 3 (Condition) x 4 (Epoch) x 7 (ROI) repeated measures ANOVA yielded a main effect of ROI, *F*(3.50, 66.57) = 22.99, *p* < .001, np2 = .548, indicating that alpha power was highest over the occipital region, lower over the temporal and frontal regions, and lowest over the central and parietal regions. There were also significant main effects of Epoch, *F*(1.37, 26.13) = 17.18, *p* < .001, np2 = .475, and Condition, *F*(2, 38) = 14.78, *p* < .001, np2 = .438. These effects were superseded by a Condition x Epoch interaction, *F*(6, 114) = 2.28, *p* = .041, np2 = .107, and a ROI x Epoch interaction, *F*(7. 41, 140.93) = 7.95, *p* < .001, np2 = .295. For the Condition x Epoch interaction, pairwise comparisons revealed alpha power across all regions to be lower for the task-related condition compared to the task-unrelated condition at *t^−2^* (*p* = .017) and *t^−1^* (*p* = .031), and lower for the task-related condition compared to both the control and task-unrelated conditions at *t^+1^* (*ps* = .016) and *t^+2^* (*ps* = .043). For the ROI x Epoch interaction, pairwise comparisons revealed all regions to significantly decrease in alpha power from *t^+1^* to *t^+2^* (rate of change (µV²): LT = −1.22, Fr =−3.21, LC = −3.06, RC = −2.56, Pa = −1.77, RT = −1.44, Oc = −2.08). However, for the frontal (*p* < .001), central (*ps* = .001), parietal (*ps* = .004), and occipital regions (*ps* = .073), this significant decrease was also evident from *t^−2^, t^−1^* and *t^+1^* relative to *t^+2^.* A marginally significant decrease in parietal alpha power was also observed from *t^−2^* to *t^+1^* (p = .08). All other interactions were non-significant (Figure 3).

**Figure 3.**
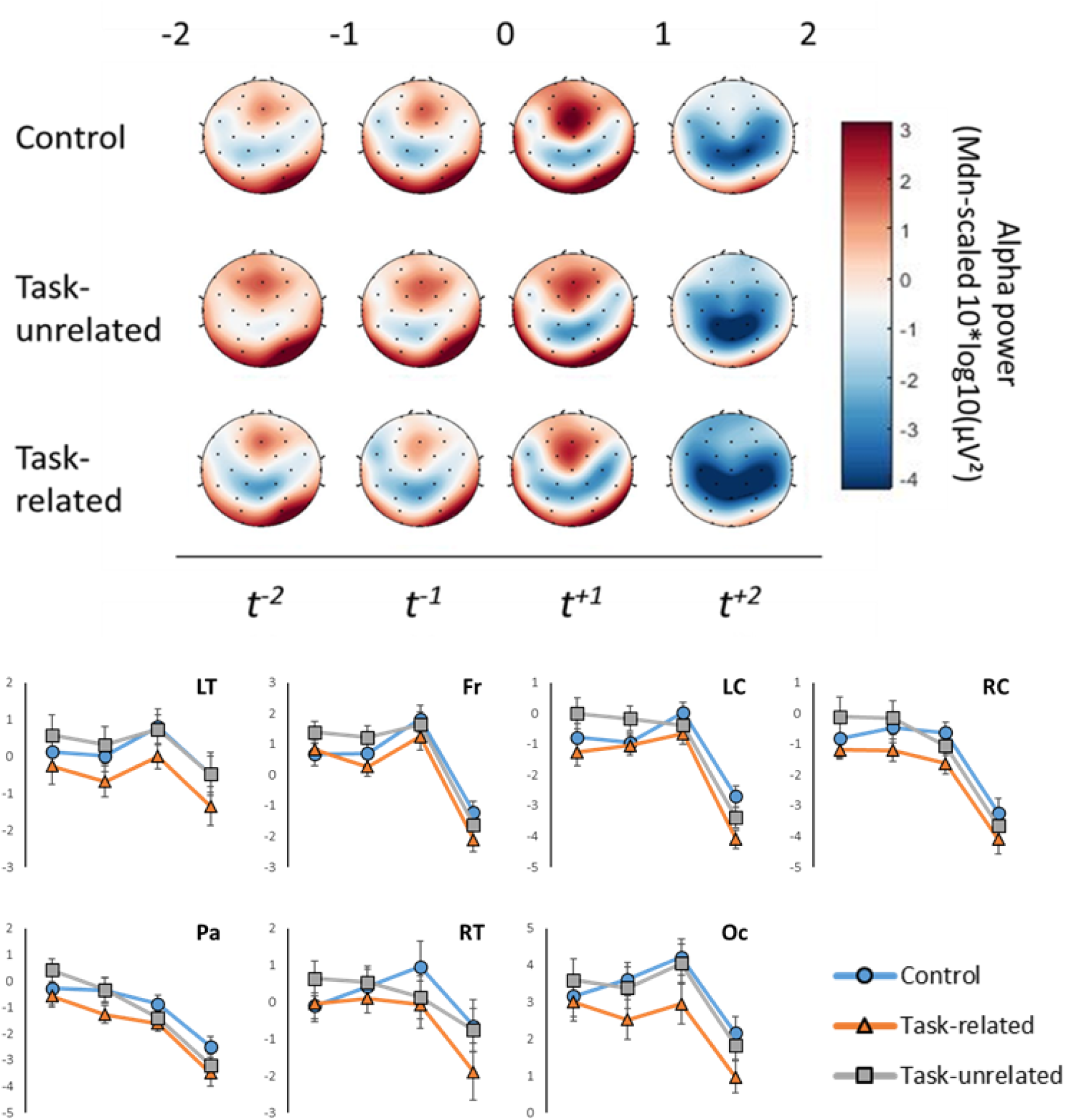
Scalp topographies (top) displaying alpha power across Epoch (*t^−2^, t^−1^, t^+1^, t^+2^;* 0 = “go” signal) for each experimental condition. Line plots (bottom) display the mean (± s.e.m) alpha power for each region of interest (left temporal = LT, frontal = Fr, left central = LC, right central = RC, Parietal = Pa, right temporal = RT, occipital = Oc) as a function of Epoch.

**Figure 4.**
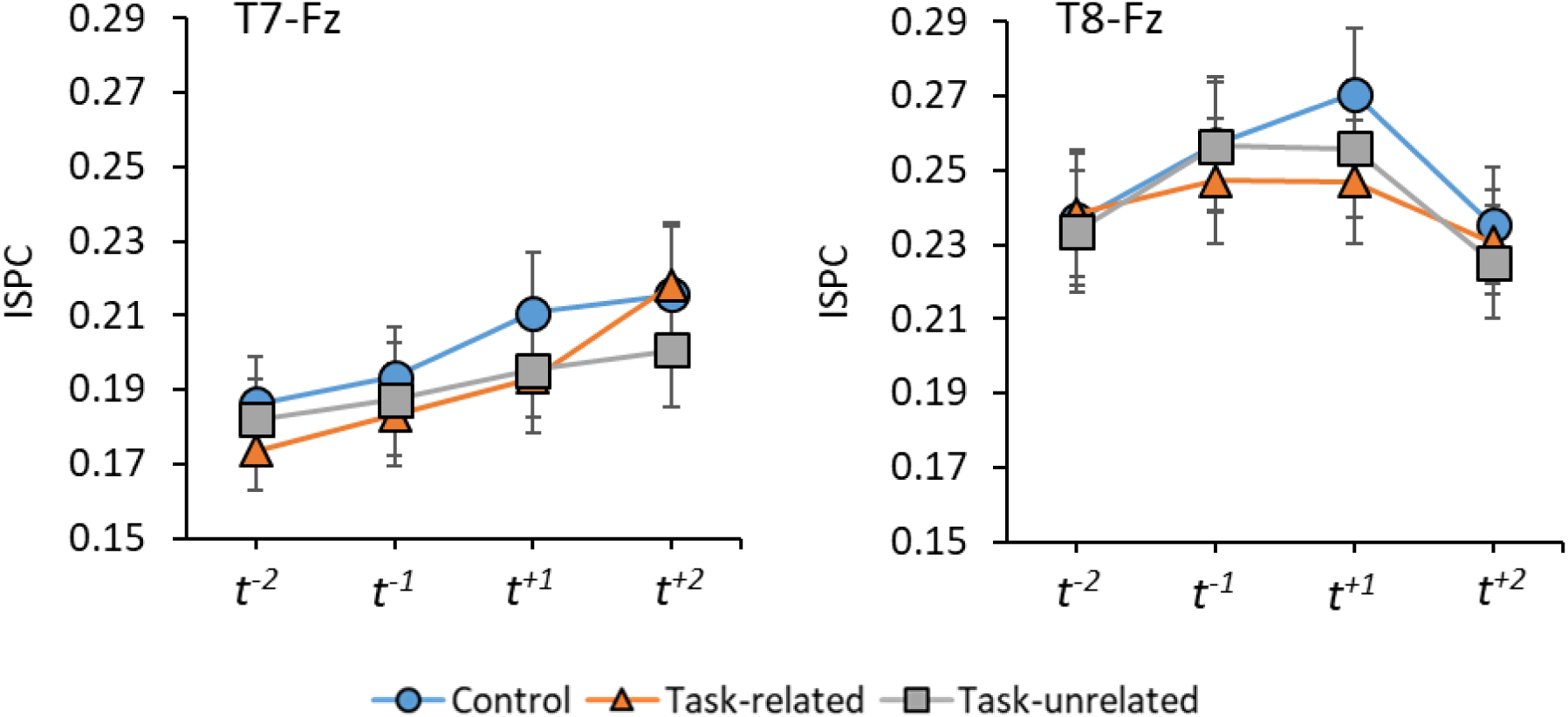
Line plots displaying the mean (± s.e.m) inter site phase clustering (ISPC) as a function of condition and epoch for both T7-Fz (left) and T8-Fz (right).

### 2.11 Alpha connectivity

A 2 (Hemisphere) x 3 (Condition) x 4 (Epoch) repeated measures ANOVA yielded a significant main effect of Hemisphere, *F*(1, 19) = 9.581, *p* = .006, np2 = .335, a marginally significant main effect of Epoch, *F*(1.58, 30.08) = 2.933, *p* = .079, np2 = .134, but no effect of Condition, *F*(1.39, 26.56) = 1.938, *p* = .173, np2 = .093. These effects were superseded by a significant Hemisphere x Epoch interaction, *F*(1.89, 35.81) = 4.764, *p* = .016, np2 = .200. Post-hoc pairwise comparisons revealed that, for all conditions, T8-Fz connectivity was significantly higher than T7-Fz connectivity at *t^−2^* (*p* = .007), *t^−1^* (*p* = .003), and *t^+1^* (*p* = .005), but not *t^+2^* (*p* = .248). They also revealed that T7-Fz connectivity increased from *t^−2^* to *t^+2^* (*p* = .072), and that T8-Fz connectivity increased from *t^−2^* to *t^−1^* (*p* = .077), and then decreased from *t^+1^* to *t^+2^* (*p* = .010).

## 3. Discussion

The present study attempted to manipulate self-talk to evaluate the fidelity of associating EEG alpha oscillations recorded from left temporal sites during motor preparation and execution with verbal-analytic processes. Our results show that both self-talk conditions (task-related and task-unrelated) were rated higher for self-talk frequency and intensity compared to the control condition, confirming that our manipulations increased verbal processing as intended. Importantly, the task-related condition was rated as being the most difficult, mentally demanding, and consciously controlled condition. Participants also performed 60% slower and decreased the speed at which they accelerated the hand when reaching for (30% slower) and transporting (40% slower) the jar. The task-related condition therefore increased verbal processing and encouraged a more effortful and conscious mode of motor control. The task-unrelated condition was rated as being more difficult than the control condition, reflected by the slower movement times (~13%). However, as there was no difference in reported levels of conscious control and mental effort, and no difference in hand acceleration profiles between the task-unrelated and control conditions, it can be argued that task-unrelated self-talk increased verbal processing without inducing increased levels of conscious movement control.

Despite both self-talk conditions increasing verbal processing compared to the control condition, our EEG data showed activity in the left-temporal region of the cerebral cortex to only increase (i.e., alpha power decreased) for the task-related condition during movement execution (*t*^+1^ and *t*^+2^). As no differences were observed between the control and self-talk conditions during movement preparation (*t*^−2^ and *t*^−1^), when participants were motionless, this question the fidelity of inferring verbal-analytic processing demands from left temporal EEG alpha power during movement preparation. Rather, our findings suggest that left temporal alpha power is likely to reflect a broader range of processes associated with conscious motor processing during movement, beyond that of language (Poldrack, 2006).

An alternative interpretation for the increased left temporal activity during task-related self-talk can be considered when acknowledging that this increase was evident across the entire scalp topography (Figure 3). Such a finding mirrors previous research that has shown less-skilled performers exhibit globally lower EEG alpha power (increased activity) compared to their higher-skilled counterparts (Del Percio et al., 2009; Parr et al., 2019). This is thought to reflect reductions in neural efficiency as individuals exert increased cognitive effort to meet the demands of the task. Indeed, alpha rhythms are proposed to reflect thalamo-cortical and cortico-cortico loops that facilitate/inhibit the transmission and retrieval of both sensorimotor and cognitive information into the brain (Deeny et al., 2003; Pfurtscheller & Lopes da Silva, 1999). Accordingly, the global decrease in alpha power during the more consciously performed task-related self-talk condition may therefore reflect the greater demands on attentional resources to process sensory-motor information as the mechanics of movements are monitored and updated online. It is important to acknowledge that the task-related self-talk condition significantly altered movement kinematics. As such, it is difficult to untangle the extent to which the present effect is driven by changes in neural efficiency, movement efficiency, or an interaction of these factors. However, such a distinction is perhaps not necessary or possible during dynamic motor tasks, given their shared association with the effortful and conscious motor control that characterise the early stages of learning (Fitts & Posner, 1967).

Our results showed no difference in T7-Fz or T8-Fz connectivity between the two self-talk conditions, despite the reported differences in verbal processing and conscious control. In fact, even with a reanalysis of these data with a spatial filter (surface Laplacian), we found the lowest T7-Fz connectivity for the more consciously performed task-related condition during jar-transportation (Appendix S3). Whilst evidence exists to suggest that increased T7-Fz connectivity is characteristic of less-skilled and more explicit motor performance (Gallicchio, Cooke, et al., 2016; Parr et al., 2019; Zhu, Poolton, Wilson, Hu, et al., 2011), the results from the present study challenge the interpretation of this measure as representing the functional communication between verbal-analytic and motor planning processes.

The absence of change in T7-Fz connectivity for the task-unrelated self-talk condition inadvertently addresses concerns that previous research utilising this measure may be confounded by between and within participant variability in the use of motivational self-talk (Bellomo, Cooke, & Hardy, 2018). Our findings suggest that changes in T7-Fz connectivity reflect processes that are not directly related to verbal processes, be it task-related or task-unrelated. However, we cannot rule out a floor effect due to participants being considered highly experienced in this rather rudimentary reaching task. In other words, it is possible that our task was not novel or difficult enough to induce the non-essential cortico-cortical communication between task-relevant and task-irrelevant cortical regions. Some support for this thesis can be taken from Parr et al. (2019), who found group-level (implicit vs explicit training) connectivity differences in a reaching and grasping task that was similar to the one employed here but which required the use of a prosthetic hand.

T7-Fz connectivity may be more influenced by aspects related to the nature of movement rather than simply verbal processing and/or conscious control. For example, we found a significant increase in T7-Fz connectivity from *t^−2^* to *t^+2^* for all conditions, suggesting a greater response to the changing demands of our task across time (i.e., movement preparation versus movement execution) than to the between-condition alterations in self-talk and conscious control. It should therefore be reiterated that phase-based connectivity merely measures the phase lag consistency between signals recorded from two sites, with relations drawn to communication pathways being inferred rather than directly assessed (Bellomo et al., 2018; Cohen, 2014). As such, it also becomes difficult to interpret our finding that T8-Fz connectivity was significantly higher than T7-Fz connectivity for all conditions across the majority of our task phases. Previously, researchers have inferred the involvement of visuospatial processes in motor performance from T8-Fz connectivity (Bellomo et al., 2018; Cooke, 2013; van Duijn, Buszard, Hoskens, & Masters, 2017). It could therefore be argued that this observed hemispheric asymmetry reflects the obvious visuospatial component of our reaching and grasping task. However, attempts to directly assess the relationship between T8-Fz upper-alpha connectivity and visuospatial processing during motor performance are required to investigate these claims, similar to how we examined the involvement of T7-Fz connectivity with verbal processing in this study (Poldrack, 2006). It could also be argued that these hemispheric differences may be driven by the right-handed nature of our task, given activation of the motor cortex usually shows contralateral bias during upper-limb control (Halsband & Lange, 2006; Kim et al., 1993). Whilst this is possible, future work is needed to explore the independent contribution of task laterality to measures of EEG cortico-cortical connectivity.

In light of our findings, several limitations should be considered. First, it could be argued that our results are limited by administering our control condition in a fixed (always first) order. However, we argue this was necessary to optimise the natural characteristics of initial performance and avoid unintentionally inducing more conscious performance and the accrual of declarative task knowledge. Second, the degree of temporal “jitter” evident in our EEG analyses should be noted. To elaborate, our data were segmented into pre-defined epoch lengths relative to the onset of movement (−2 to +2 seconds) to enable meaningful comparisons to be made between conditions. However, this method fails to guarantee the segmented data represent the same phase of movement (from 0 to +2 seconds) on a trial-to-trial basis, especially given the differences in movement time. Although less than ideal, our measures of alpha power and connectivity were highly consistent across time for all conditions, suggesting a minimal effect of such temporal lag. Finally, our analyses are potentially limited by solely examining alpha-based connectivity to infer cortico-cortical networking. Indeed, it is still unclear how the interpretation of alpha connectivity differs from the interpretation of non-alpha connectivity (e.g. theta, ~4 – 8 Hz). Future research could pay greater attention to this issue and include measures of functional connectivity that either include several frequency bands or do not rely on a specific frequency band (e.g., Granger causality, Phase Slope Index).

In conclusion, our results failed to endorse EEG alpha activity recorded from the left temporal region as a valid index of verbal analytic processing demands during a motor task. Instead, our results suggest that increased left temporal alpha activity exhibited during more consciously controlled motor performance should be attributed to a spatially broader phenomenon consistent with decreased neural efficiency. Furthermore, the approach presented in this study invites motor control scientists to be cautious when inferring a certain cognitive process based solely on local activity. We encourage researchers to explore how cognition maps onto regional brain activity considering the whole topography, specifically during the performance of motor tasks to improve our understanding of how the brain controls movements.

## Appendix 1 Manipulations to ensure adherence to task-related self-talk instructions

Photos depicting the placement of electrodes both on the workspace and arm to encourage full adherence to the task-related self-talk instructions. Manipulations were designed to encourage a more conscious mode of movement control.

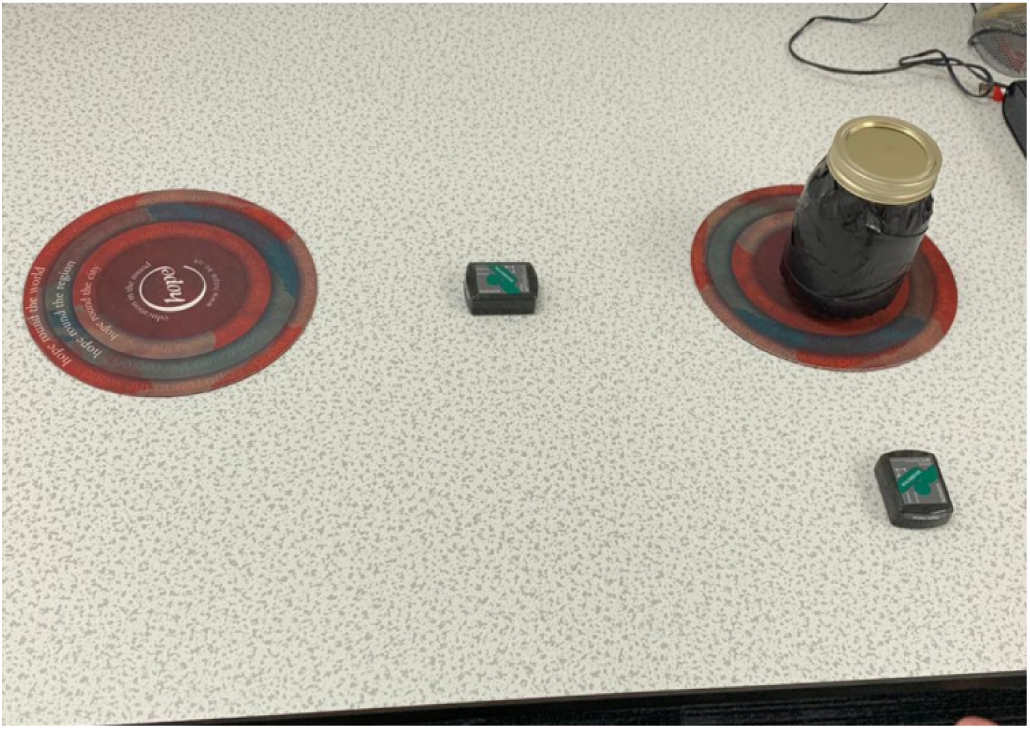
“Keep hand 5cm above table”. Two additional electrodes were placed halfway between the hand start location and the jar, and halfway between the jar and target locations. Participants were informed that the distance between the electrode on hand and the electrodes on the workspace would be recorded during performance. As such, participants were instructed to ensure this distance was as close to 5cm as possible.

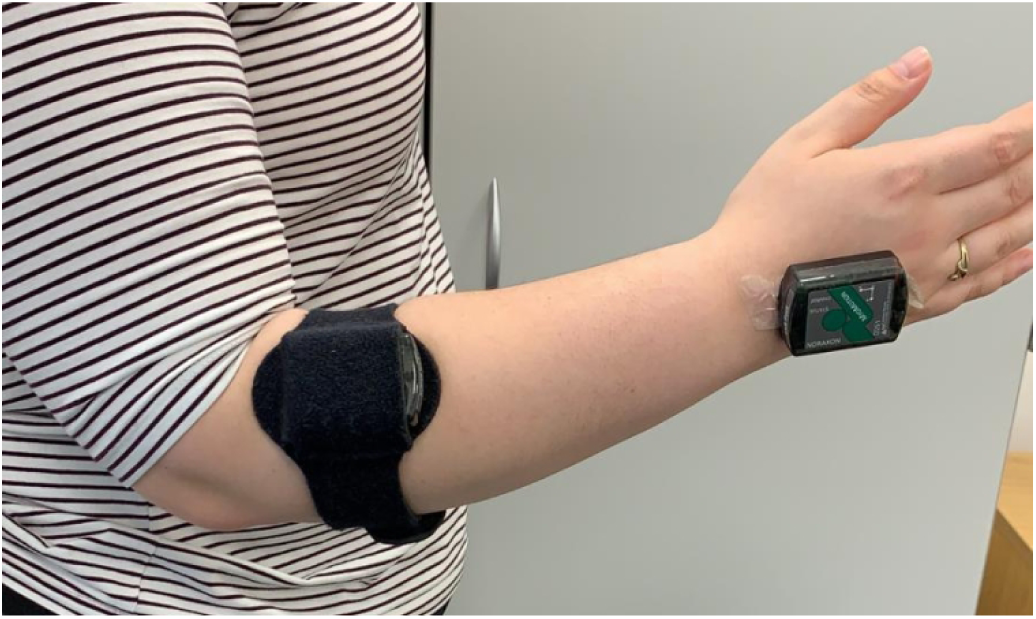
“Keep elbow below the wrist”. An additional electrode was placed towards the elbow using a Velcro strap. Participants were informed that the electrode on the elbow must remain below the electrode on the hand throughout task performance. Participants were told this would be objectively assessed.

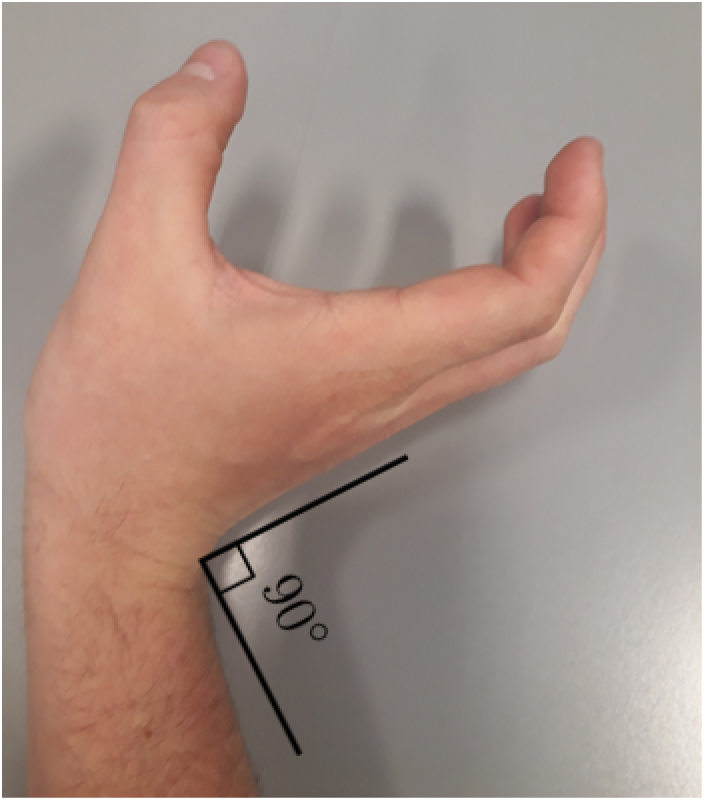
“Keep wrist flexed at 90 degrees”. Participants were required to perform the task with their hand hyperextended to create a 90 degree angle with the forearm (pictured). Participants were told that their adherence could be objectively monitored using the electrode placed on the hand (not pictured).

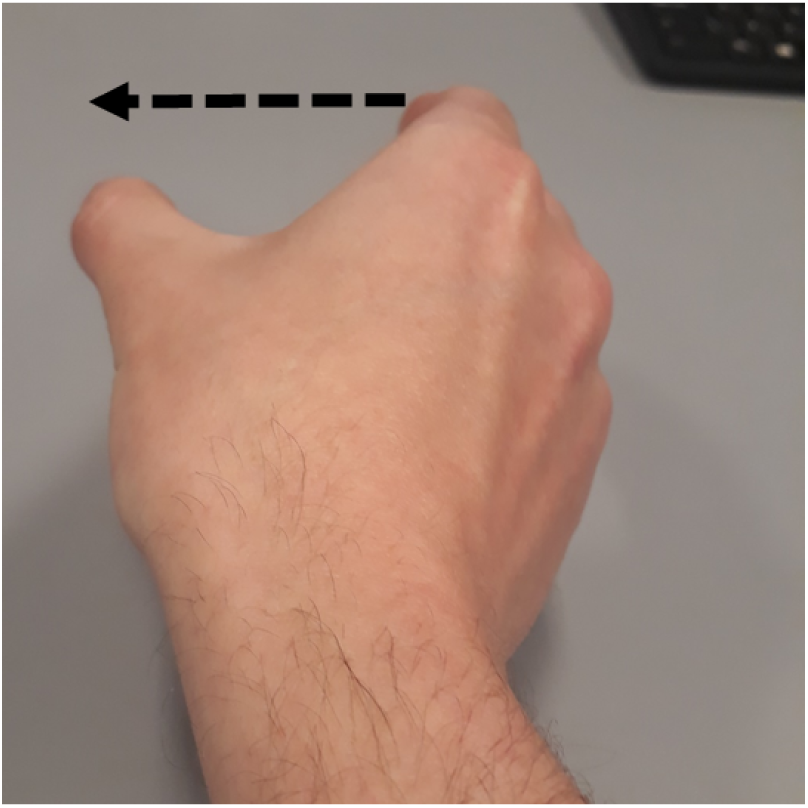
“Keep thumb below the index”. Participants were required to rotate their hand in a manner that ensure the thumb was always positioned lower than the index finger (as opposed to being in-line. Again, participants were told that adherence would be objectively assessed by calculating the angle of the hand electrode during performance (not pictured).

## Appendix 2 Individual alpha frequency identification

As the actual alpha frequency band can show inter-subject variability, we attempted to specify each participant’s individual alpha frequency band (IAF). The below figure displays the mean spectra recorded over occipital-parietal electrodes during a 1-minute eyes closed resting baseline. For the line plots that are in red, IAF was taken as the peak frequency occurring between 7 and 15 Hz, as a clear single peak could be observed upon inspection within this band. For the line plots that are in black, a clear single peak was not visible, resulting in the IAF being taken as the power spectra weighted mean (centre of gravity) within the 7 – 15 Hz band. For one participant, baseline data was corrupt resulting in IAF being set at the typical 10 Hz.

**Fig 1.**
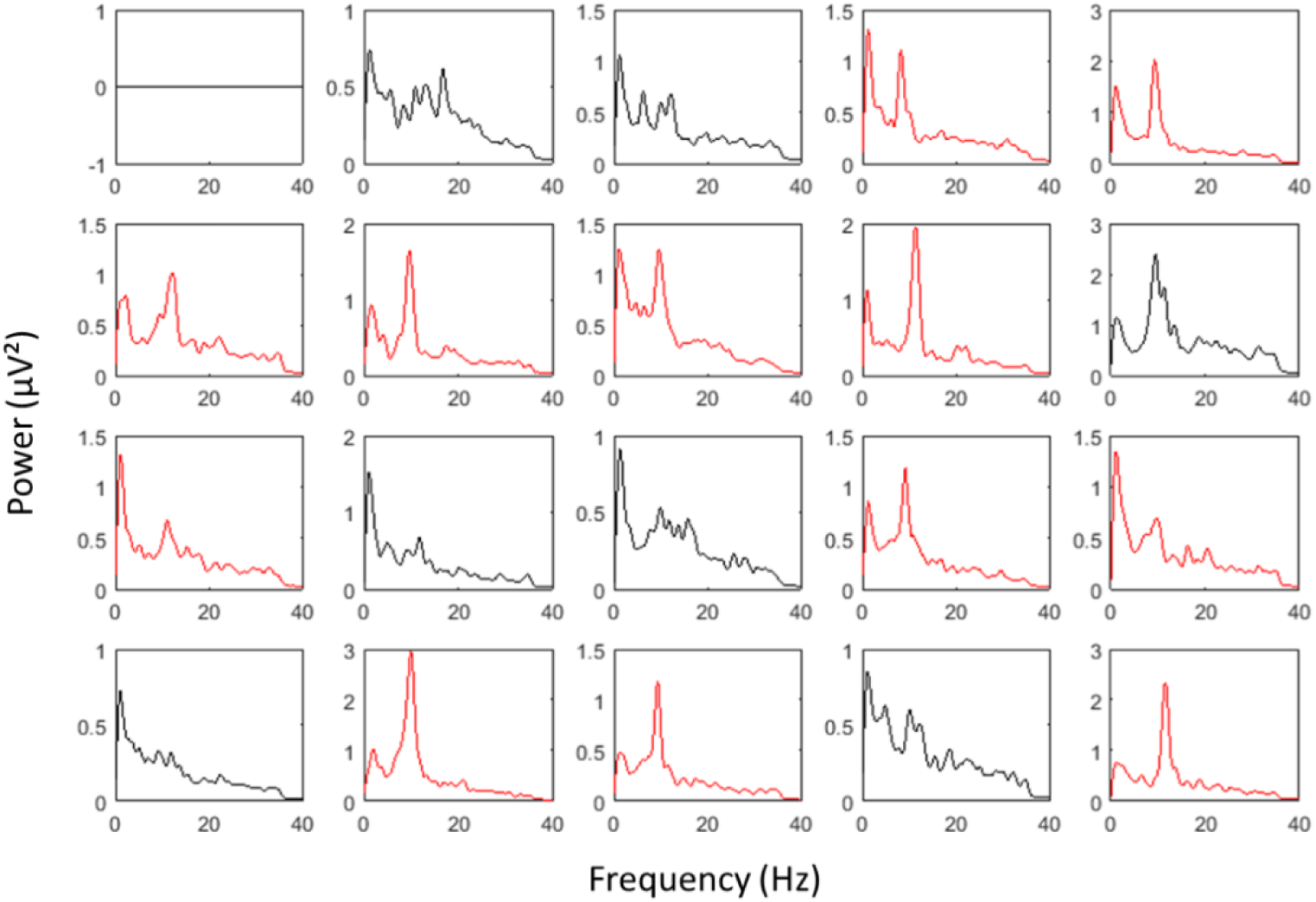
Line plots displaying mean spectral power recorded over occipital-parietal electrodes during a 1-minute eyes closed resting baseline. Each line plot represents a single participant. Red lines indicate the identification of a single peak alpha frequency, whilst black lines indicate the calculation of the centre of gravity.

## Appendix 3 Spatially enhanced analyses of alpha power and connectivity using surface Laplacian and channel levels analyses

To enhance the local features of our data, we applied a scalp-level surface Laplacian transformation (Cohen, 2014; function available at http://mikexcohen.com/lectures.html), which acts as a spatial band-pass filter to attenuate the effects of volume conduction. In line with our aim to highlight local features, we conducted analyses for alpha power on a channel level over sites T7, T8, Fz, CP1, CP2, FC1, FC2, O1, O2. These channels were chosen based upon previous literature and topographical inspection of our data (Figure 1).

**Figure 1.**
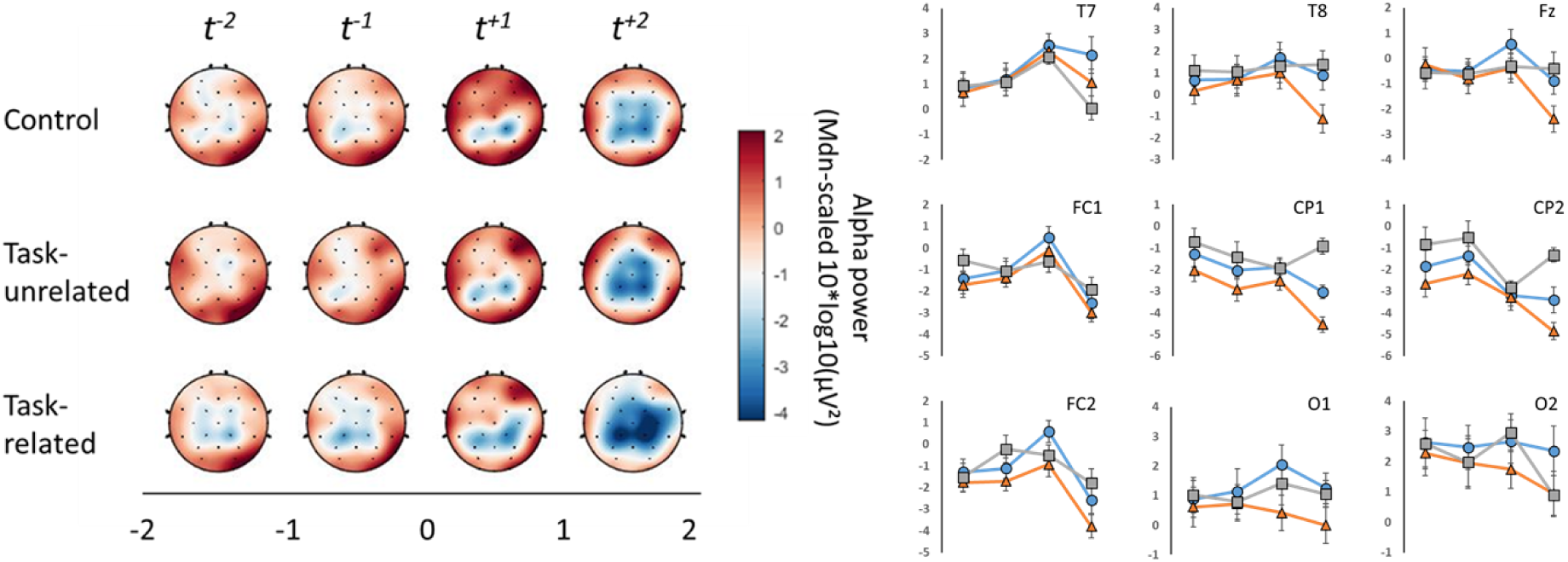
Scalp topographies (left) and line graphs (right) depicting surface Laplacian spatially enhanced regional alpha power across time, relative to movement onset, for each experimental condition.

### Results

#### Alpha power

Results from a 3 (Condition) x 4 (Epoch) x 9 (Channel) repeated measures ANOVA showed a significant main effect of Condition, *F*(2, 38) = 11.168, *p* < .001, np2 = .370, a main effect of Epoch, *F*(1.36, 25.85) = 6.44, *p* = .011, np2 = .253, and a main effect of channel, *F*(3.74, 71.13) = 16.103, *p* < .001, np2 = .459. These effects were superseded by significant Condition x Epoch, *F*(6, 114) = 3.606, *p* = .003, np2 = .160, and Epoch x Channel, *F*(24, 456) = 7.126, *p* < .001, np2 = .273, interactions. Due to the aim of this analysis, we only further examined effects involving the factor Channel. For the Epoch x Channel interaction, Bonferroni pairwise comparisons revealed that whilst no channels differed in activity from *t^−2^* to *t^−1^*, the onset of movement from *t^−1^* to *t^+1^* saw a significant increase in alpha power at T7 (*p* = .041) and a significant decrease in alpha power at CP2 (*p* = .024) and FC1 (*p* = .018). The transition from *t^+1^* to *t^+2^* then saw significant reductions in alpha power at T8 (*p* = .013), Fz (*p* = .003), CP1 (*p* = .001), CP2 (*p* = .009), FC1 (*p* < .001) and FC2 (*p* < .001). Channels O1 and O2 did not change across the entirety of the task.

#### Alpha connectivity

A 3 (Condition) x 2 (Hemisphere) x 4 (Epoch) repeated measures ANOVA showed no main effect of Hemisphere, *F*(1, 19) = 2.339, *p* = 1.43, np2 = .110, or Epoch, *F*(3, 57) = 0.831, *p* = .482, np2 = .042. There was however a significant main effect of Condition, *F*(2, 38) = 4.048, *p* = .025, np2 = .176, that was superseded by a significant Epoch x Condition interaction, *F*(6, 114) = 2.489, *p* = .027, np2 = .116. Pairwise comparisons revealed that the task-related condition produced a significant decrease in temporal-frontal connectivity at *t^+2^* relative to all previous time-points (*ps* = .031), and relative to the control (*p* = .017) and task-unrelated (*p* = .02) conditions (Figure 2). All other interactions were non-significant.

**Fig 2.**
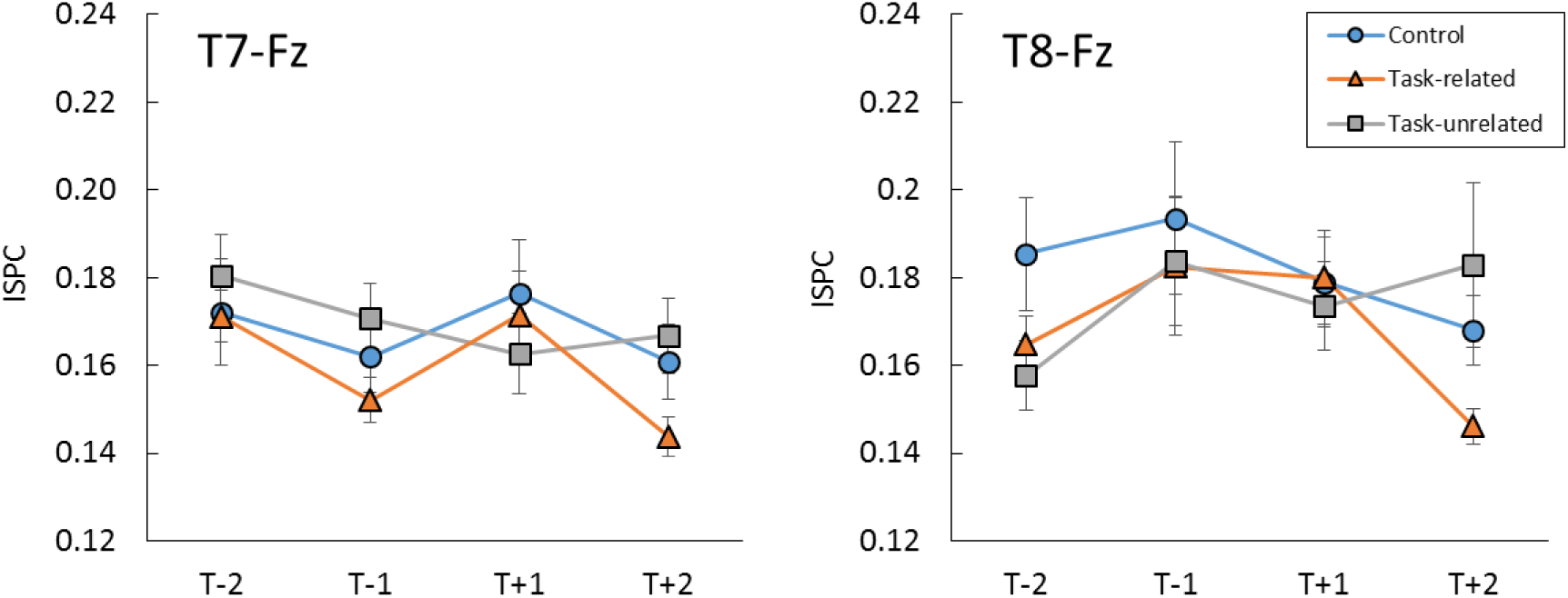
Line plots displaying the mean (± s.e.m) intersite phase clustering (ISPC) as a function of condition and epoch for both T7-Fz (left) and T8-Fz (right).

## Footnotes

1. Sites T7 and T8 are sometimes referred to as T3 and T4, respectively, in other EEG systems.

